# Cancer related subarachnoid hemorrhage: a multicenter retrospective study using propensity score matching analysis

**DOI:** 10.1101/2022.03.10.483885

**Authors:** Senthil K

## Abstract

**Objective:** To investigate the clinical features, risk factors and underlying pathogenesis of cancer related subarachnoid hemorrhage (SAH).

**Methods:** Clinical data of SAH in patients with active cancer from January 2010 to December 2020 at four centers were retrospectively reviewed. Patients with active cancer without SAH were matched to SAH patients with active cancer group. Logistic regression was applied to investigate the independent risk factors of SAH in patients with active cancer, after a 1:1 propensity score matching. A receiver operator characteristic curve was configured to calculate the optimal cut-off value of the joint predictive factor for cancer related SAH.

**Results:** A total of 82 SAH patients with active cancer and 309 patients with active cancer alone were included. Most SAH patients with cancer had poor outcomes, with thirty-day mortality of 41.5%, and with ninety-day mortality of 52.0%. The propensity score matching yielded 75 pairs of study participants. Logistic regression revealed that a decrease in platelet and prolonged prothrombin time were the independent risk factors of cancer related SAH. In addition, receiver operator characteristic curve of the joint predictive factor showed the largest AUC of 0.8131, with cut-off value equaling to 11.719, with a sensitivity of 65.3% and specificity of 89.3%.

**Conclusions:** Patients with cancer related SAH often have poor outcomes. The decrease in platelet and prolonged prothrombin time are the independent risk factors of cancer related SAH, and the joint predictive factor with cutoff value equal to 11.719 should hence serve as a novel biomarker of cancer related SAH.

## INTRODUCTION

Cancer related stroke has recently received increasing attention from clinicians. Although the risk of hemorrhagic stroke and the risk of ischemic stroke are almost equal in patients with cancer ^1^, hemorrhagic stroke seems to receive less attention than ischemic stroke. This is because clinicians often view hemorrhagic stroke as a catastrophic and terminal event^2^. As a result, data about cancer patients with hemorrhagic stroke, especially on SAH are not much. Previous study with small samples shows that SAH may occur in patients with various kind of cancer, including primary intracranial tumors, spinal tumors, somatic solid tumors, and hematological tumors. It was also found that the pathogenesis of SAH in patients with cancer is associated with intratumoral hemorrhage in intracranial primary tumors or metastatic tumor. Pathogenesis of SAH is also associated with the blood coagulation dysfunction in no intracranial cancer. It has also been reported that most of SAH in patients with cancer tend to have poor prognosis ^3-5^. In 2010, systematic retrospective clinical research on cancer with intracranial hemorrhage found that among 181 active cancer patients with intracranial hemorrhage, 48(25.4%) patients had SAH. The study revealed that intratumoral hemorrhage (61%) and coagulopathy (46%) accounted for the majority of hemorrhages, whereas hypertension (5%) was rare, with median survival of 3 months (95% confidence interval [CI] 2–4), and its 30-day mortality of 31%^6^. In 2016, a different study reported that previous cancer history was a risk factor for the poor functional outcome of SAH^7^. However, the risk factors of SAH in patients with active cancer and the relationship between active cancer and SAH have not been well elucidated.

Therefore, the present study aimed to investigate the clinical features, risk factors and underlying pathogenesis of cancer related SAH by conducting a multicenter retrospective study using propensity score matching analysis. It was hypothesized that active cancer may play an important role in the development of SAH in patients with active cancer. Eligible patients with active cancer related SAH were enrolled and age and sex matched patients with active cancer alone were also enrolled in the ratio of nearly 1:4. The univariate and multivariate binary logistic regression were applied before and after propensity score matching (PSM) to investigate independent risk factors and underlying pathogenesis of cancer related SAH. Thus, this study may contribute to establishing the relationship between SAH and the cancer. Additionally, this study may help in detecting the patients at high risk of SAH and help clinicians to take effective preventive and therapeutic measures for such patients.

## METHODS

### Patients

We retrospectively analysed the records of all SAH patients with active cancer from Guangxi medical university affiliated tumor hospital, the first, second and fourth affiliated hospitals of Guangxi medical university between January 2010 and December 2020. Patients met the following recruitment criteria were enrolled in SAH with active cancer group: (1) age ≥18 years; (2) spontaneous SAH was confirmed by CT, MRI, or lumbar puncture; (3)cancer was demonstrated by pathological examination; and (4) active cancer was defined as a diagnosis of cancer within 6 months before enrollment, any treatment for cancer within the previous 6 months, or recurrent or metastatic cancer. The exclusion criteria of SAH in patients with active cancer group included: (1) age <18 years; (2) patients had postoperative hemorrhages; (3) cancer patients with SAH due to trauma, traditional aneurysm, cerebrovascular malformations, moyamoya disease, intracranial venous thrombosis; (4) incomplete medical records. Patients with active cancer alone were matched to SAH in patients with active cancer group by tumor types were recruited at a ratio of nearly 4:1 as control group. Our records search was as outlined in Appendix 1:Figure S1.

### Data collection and follow-up

Clinical data of all patients, including age, gender, vascular risk factors, type of cancer, etiology of subarachnoid hemorrhage in patients with cancer, presentation, current cancer treatment, vascular risk factors, and outcome were noted. In addition, laboratory data at diagnosis were also recorded including routine blood tests, blood biochemistry, coagulation function and levels of plasma D-dimer as well as tumor markers. Findings of imaging examinations such as echocardiography, transcranial Doppler ultrasound(TCD), cranial CT, computed tomography angiography (CTA), MRI, magnetic resonance angiography (MRA) and digital subtraction angiography (DSA) were also collected. In the present study, the following criteria were employed to diagnose coagulopathy: platelets<100/mm^3^, prothrombin time>13 seconds, international normalized ratio >1.5, activated partial thromboplastin time>45 seconds and disseminated intravascular coagulation (DIC) (fibrinogen<200 mg/dL and D-dimer>290 ng/dL). The severity of clinical status was assessed by Hunt & Hess grade.

### Statistical analysis

Statistical analysis was performed using SPSS version 25.0 software (IBM). Primarily, each variable was tested for normal distribution using the Kolmogorov-Smirnov test. The data was then expressed as mean ± standard deviation for normally distributed variables or median (25–75%) for the variables without normal distribution and the classified variable was represented by count (percentage). The Mann-Whitney u-test was used for non-normally distributed variables; unpaired comparisons were made with the student’s unpaired t-test, while Pearson’s χ^2^ or Fisher’s exact test was used to compare categorical variables. Besides, the Kaplan–Meier method was used to estimate the rate of cumulative events, and differences between groups were assessed using the log-rank test. A Cox regression model was constructed to evaluate the hazard ratio for 30- and 90-day mortality in cancer with SAH group and predictors of mortality univariate. Variables with *P*<0.05 in univariate analyses were considered explanatory variables and were entered to multivariate models with entry and exit levels set at 0.05 to produce the final multivariate analyses.

To eliminate the impacts of conventional vascular risks on subarachnoid hemorrhage in cancer patients, a propensity score matching (PSM), using a multivariable logistic regression model based on: age, gender, previous strokes, coronary heart disease, hypertension, diabetes, current smoker, and drinking was performed. As a result, pairs of patients receiving cancer with SAH group or active cancer group alone were derived using 1:1 greedy nearest neighbor matching within propensity score of 0.02. This strategy resulted in 75 matched pairs in each group. The balance of variables between groups after propensity score-matching was analyzed using a paired *t*-test for continuous measures and the McNemar test (for categorical variables). Moreover, for nonnormal data, the Wilcoxon sum rank test was used (Wilcoxon signed rank test for paired within-group comparisons).

To find independent risk factors of SAH in patients with cancer, univariate and multivariate binary logistic regression was applied. Univariate significant factors at *P<0*.*05* were included in the stepwise logistic regression model, with entry and exit levels set at 0.05 for the final multivariate analysis. Receiver operating characteristic (ROC) curve analysis was used to evaluate the diagnostic performance of the resulting regression model based on its AUC, sensitivity, and specificity values. Optimal cut-off value was determined by the Youden index. All *p-*Values were two-sided, and *P*<0.05 was considered statistically significant. No formal calculation of sample size was performed because of the characteristics of cancer with SAH.

### Ethics

Ethical approval of this study was provided by the Guangxi Medical University Review Board. The written informed consent was waivered because of the retrospective nature of our study.

## RESULTS

### Patient profiles

Among the 859 cancer patients with intracranial hemorrhage(ICH) admitted to the four centers during the study period, 82 active cancer patients with SAH meeting all eligibility criteria and were included in the final analysis. Concurrently, regarding the control group, 309 patients with active cancer alone were matched to the active cancer patients with SAH by tumor types at a ratio of nearly 4:1. Moreover, after propensity matching, a total of 150 patients were included in the study population. The propensity score matching yielded 75 pairs of study participants in an unbiased database (Table 1). Before the propensity score-matching was performed, cancer patients with SAH group contained 56 (68.3%) men and 26 (31.7%) women. The median age was 53 (range 44 –66) years.

**Table 1.**
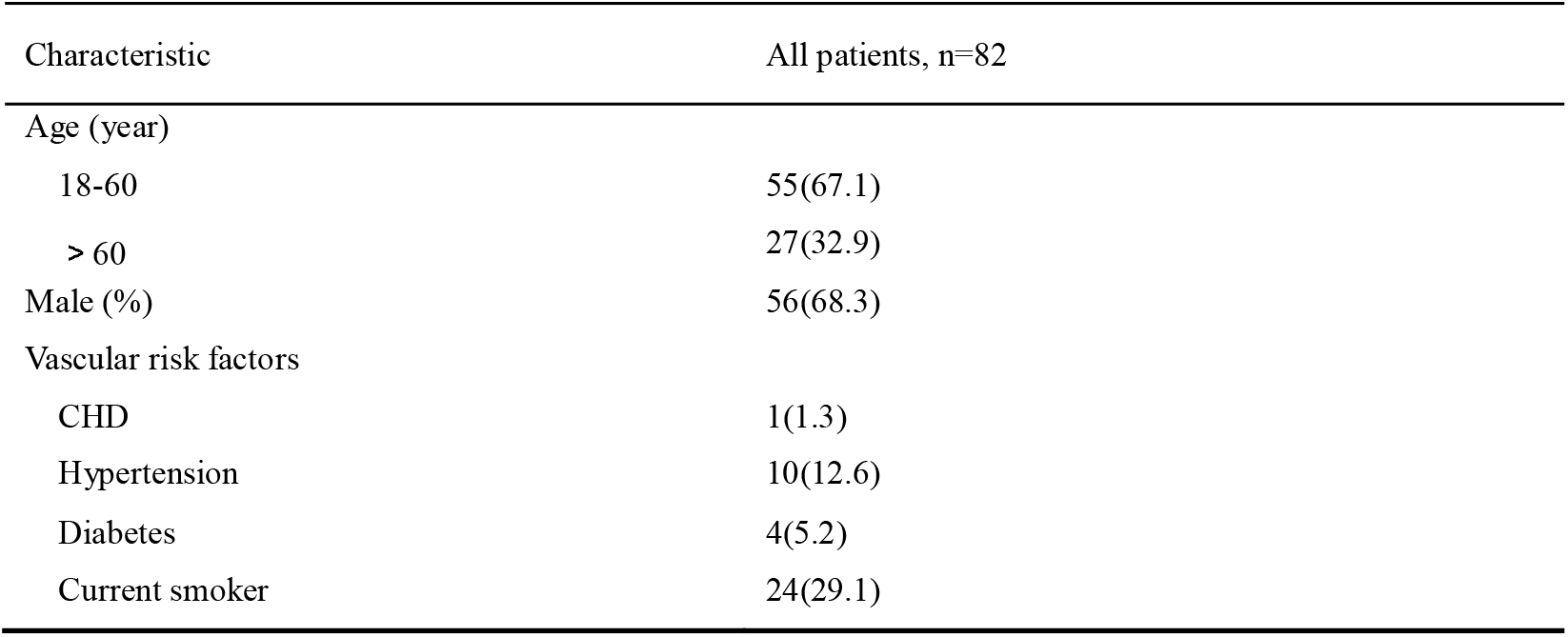
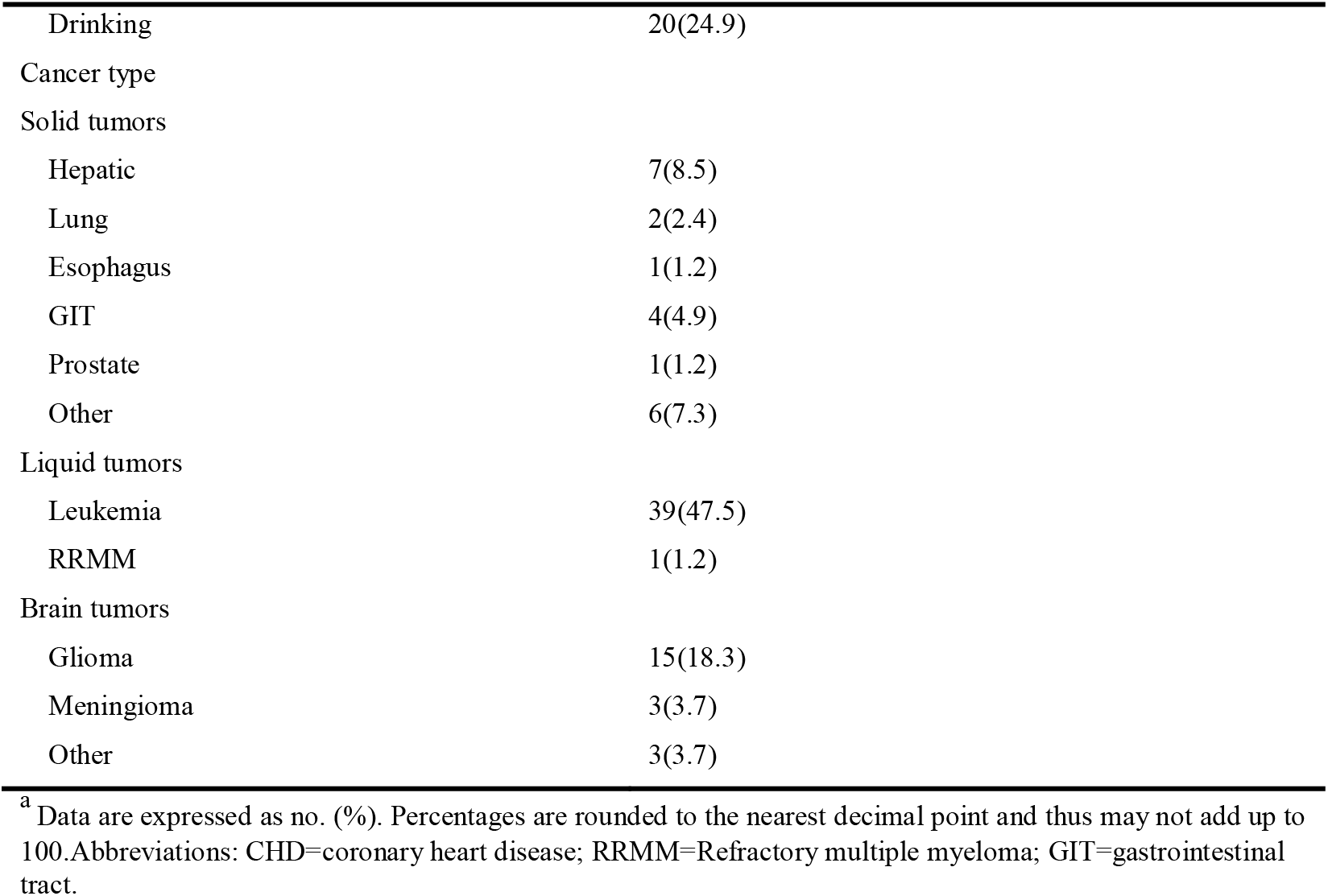
Demographics of cancer patients with subarachnoid hemorrhage ^a^

| Characteristic | All patients, n=82 |
| --- | --- |
| Age (year) |  |
| 18-60 | 55(67.1) |
| > 60 | 27(32.9) |
| Male (%) | 56(68.3) |
| Vascular risk factors |  |
| CHD | 1(1.3) |
| Hypertension | 10(12.6) |
| Diabetes | 4(5.2) |
| Current smoker | 24(29.1) |
| Drinking | 20(24.9) |
| Cancer type |  |
| Solid tumors |  |
| Hepatic | 7(8.5) |
| Lung | 2(2.4) |
| Esophagus | 1(1.2) |
| GIT | 4(4.9) |
| Prostate | 1(1.2) |
| Other | 6(7.3) |
| Liquid tumors |  |
| Leukemia | 39(47.5) |
| RRMM | 1(1.2) |
| Brain tumors |  |
| Glioma | 15(18.3) |
| Meningioma | 3(3.7) |
| Other | 3(3.7) |

<sup>a</sup> Data are expressed as no. (%). Percentages are rounded to the nearest decimal point and thus may not add up to 100. Abbreviations: CHD=coronary heart disease; RRMM=Refractory multiple myeloma; GIT=gastrointestinal tract.

### Cancer type

The cancer contributing to SAH was a solid tumor in 21 patients (25.6%), a primary brain tumor in 21 (25.6%) and a hematologic tumor in 40 (48.8%). The most common pathological type of cancer was Leukemia (47.5%), followed by glioma (18.3%) and hepatoma (8.5%). After diagnosis of cancer, 47 (57.3%), 10 (12.2%), and 12 (14.6%) patients experienced SAH in the first 6 months, 6 months to 1 year and >1 year, respectively. Additionally, 13 (15.9%) cancer patients have SAH as the first presentation (Appendix 2:Figure S2 and Appendix 5: Table S1).

### Presentation and etiology

The presentation and pathogenesis of SAH in cancer group was as summarized in Table 2. Patients with active cancer and SAH present with the usual symptoms of SAH such as headache and vomiting. However, the SAH in patients with active cancer showed unique pathogenesis. In the present study, it was found that the possible etiology of SAH was mostly related to coagulopathy (52.4%), followed by intratumoral hemorrhage (ITH) (9.6%), and was rarely related to coagulopathy combined with ITH. However, the pathogenesis of SAH in 27 (33.0%) patients was found to be indefinite. For patients with liquid tumors such as Leukemia, SAH was mostly due to coagulation dysfunction, whereas for patients with brain tumors, SAH was commonly caused by intratumoral hemorrhage.

**Table 2:** Presentation and etiology of subarachnoid hemorrhage in patients with cancer ^a^

| Characteristic | All patients,<br>n=82 n(%) | Solid tumor,<br>n=21 n(%) | Liquid tumor,<br>n=40 n(%) | Brain tumor,<br>n=21 n(%) |
| --- | --- | --- | --- | --- |
| Presentation |  |  |  |  |
| Disturbance of consciousness | 45(54.9) | 11(52.3) | 23(57.5) | 11(52.4) |
| Hydrocephalus | 8(9.8) | 2(9.5) | 1(2.5) | 5(23.8) |
| Coma | 35(42.7) | 11(52.3) | 16(40.0) | 8(38.1) |
| Seizure | 9(11.0) | 1(4.8) | 3(7.5) | 5(23.8) |
| Hemiparesis | 18(22.0) | 6(28.6) | 7(17.5) | 5(23.8) |
| Aphasia | 10(12.2) | 3(14.3) | 6(15) | 1(4.8) |
| Dysesthesia | 5(6.1) | 1(4.8) | 1(2.5) | 3(14.3) |
| Headache | 57(69.5) | 16(76.2) | 29(72.5) | 12(57.1) |
| Nausea or vomiting | 35(42.7) | 9(42.9) | 20(50) | 6(28.6) |
| Etiology |  |  |  |  |
| ITH | 8(9.6) | 1(4.8) | 1(2.5) | 6(28.6) |
| Coagulopathy | 43(52.4) | 7(33.3) | 34(85) | 2(9.5) |
| ITH and coagulopathy | 4(4.9) | 2(9.5) | 0(0.0) | 2(9.5) |
| Cryptogenic | 27(33.0) | 11(52.3) | 5(12.5) | 11(52.3) |

<sup>a</sup> Data are expressed as no. (%). Percentages are rounded to the nearest decimal point and thus may not add up to
100. Abbreviations: ITH=intratumoral hemorrhage

### Imaging features

Radiologic investigation with computed tomography of the brain and computed tomography angiography revealed typical manifestations of cancer with SAH (Figure 1 A-F).

**Figure 1.** Imaging features: Figure 3A, 3B, and 3C were from a 28-year- female with astrocytic glioma experienced SAH due to intratumoral hemorrhage in the first 11 days after the diagnosis of astrocytic glioma. It shows a tumor in the brain (white triangle) and subarachnoid hemorrhage (thin white arrow) in Figure 3A, intratumoral bleeding in Figure 3B (thick white arrow), normal cerebral arteries in Figure 3C. Figure 3D, 3E and 3F were from a middle-aged female patient with acute monocytic leukemia combined with coagulopathy suffered from SAH. Figure 3D and 3E showed massive subarachnoid hemorrhage with ventricle hematoma. Figure 3F shows normal cerebral arteries.

### Severity score of SAH

Results of the severity score of cancer patients with SAH were as shown in Appendix 3:Figure S3. Generally, twelve (16 %) cancer patients with SAH were found with 5 scores of Hunt & Hess grade.

### Treatment

In the present study, SAH in patients with active cancer received stander treatment for SAH and cancer according to guidelines for SAH and various types of cancer.

### Survival analysis

Survival analysis using Kaplan–Meier curves showed that most SAH patients with cancer had poor outcomes, with thirty-day mortality 41.5%, and with ninety-day mortality 52.0%. Moreover, the survival time of cancer patients with SAH differed (*p* =0.041) depending on the type of cancer, with median survival of 22 days for patients with liquid tumors and 27 days for patients with solid tumors. However, due to the relatively long survival time of brain tumors and the limitation of follow-up time, the median survival data for the time could not be obtained (Figure 2A). Furthermore, the survival time of patients with cancer related SAH based on etiology showed difference (*p* =0.038), with median survival of 21 days for patients with intratumoral hemorrhage, 19 days for patients with coagulopathy and 8 days for patients with both intratumoral hemorrhage and coagulopathy (Figure 2B).

**Figure 2.** Survival curves by cancer type and etiology of SAH. (A) Cancers were categorized as solid, liquid, or brain tumor. Kaplan–Meier curves showed survival time of cancer patients with SAH differed (p=0.041) by cancer type. (B) Etiology of SAH was grouped as intratumoral hemorrhage, coagulopathy, both intratumoral hemorrhage and coagulopathy, or other diagnoses. Survival analysis based on the etiology of SAH showed survival time difference (p=0.038). Abbreviations: ITH=intratumoral hemorrhage.

Survival analysis of Cox regression model showed that 7 variables were significant predictors of 30-day mortality: having coagulopathy, aphasia, activated partial thromboplastin time, gamma-glutamyl transferase, atrial fibrillation, prealbumin and cryptogenic SAH. On the other hand, 5 variables were found to be significant predictors of 90-day mortality: having liquid tumor, activated partial thromboplastin time, prealbumin, cryptogenic SAH and coagulopathy. The results of univariate analysis of continuous and categorical variables relating to 30-day and 90-day mortality survival rate in original 82 cancer patients with SAH were as presented in Appendix 6: Table S2, Appendix 7: Table S3, Appendix 8: Table S4, and Appendix 9: Table S5. In the multivariate Cox regression models, the predictors of 30-and 90-day mortality in the whole cohort were summarized in Figure 3 and Appendix 10: Table S6.

**Figure 3.** Predictors of mortality via Cox regression models. (A) Predictors of 30-day mortality via Cox regression models. (B) Predictors of 90-day mortality via Cox regression models. HR>1 implies unfavorable prognosis for SAH in patients with active cancer. HR of 1 corresponds to no effect. Abbreviations: HR= hazard ratio; 95% CI=95% confidence interval; SAH=subarachnoid hemorrhage; APTT= activated partial thromboplastin time; PAB=prealbumin; GGT=gamma-glutamyl transferase; AF= Atrial fibrillation.

### Univariate analysis and multivariate analysis

Before propensity score matching was performed, a univariate analysis between the 82 SAH in patients with active cancer and 309 patients with active cancer alone showed that nineteen variables were significantly related to the onset of SAH. The variables were: sex; having a previous stroke; hypertension; smoker; drinking; having a brain tumor; the level of white blood cell, red blood cell, hemoglobin, platelet, neutrophil percentage, lymphocyte, lymphocyte percentage, total bilirubin, gamma-glutamyl transferase, prealbumin, uric acid, prothrombin time and international normalized ratio (Table 3)

**Table 3.**
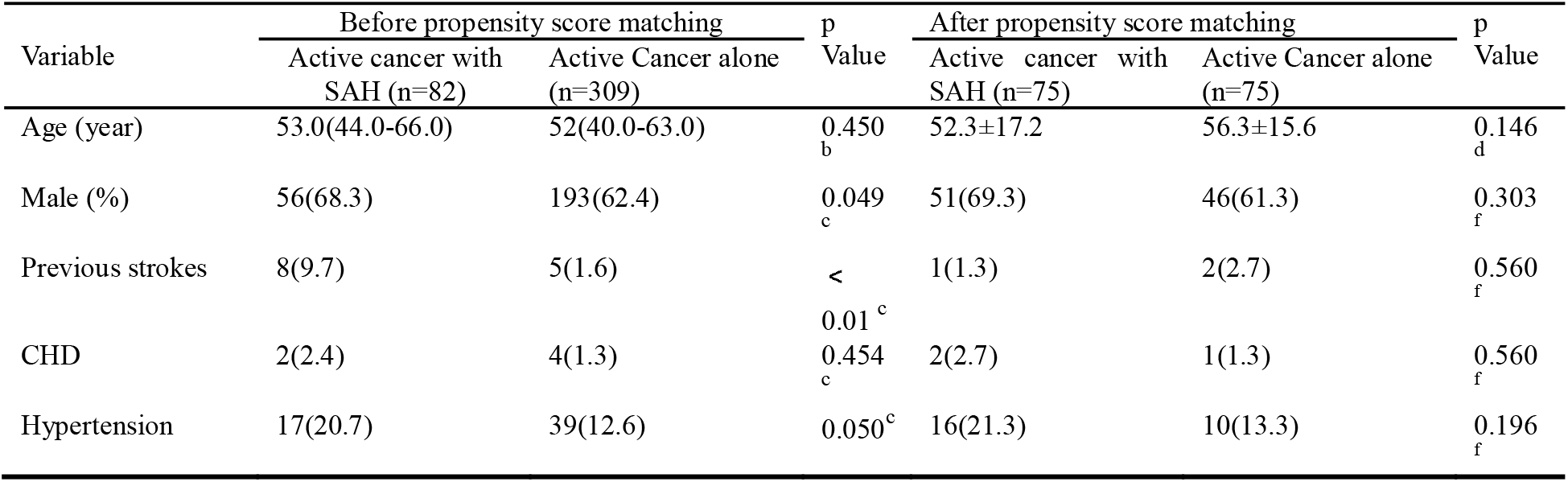
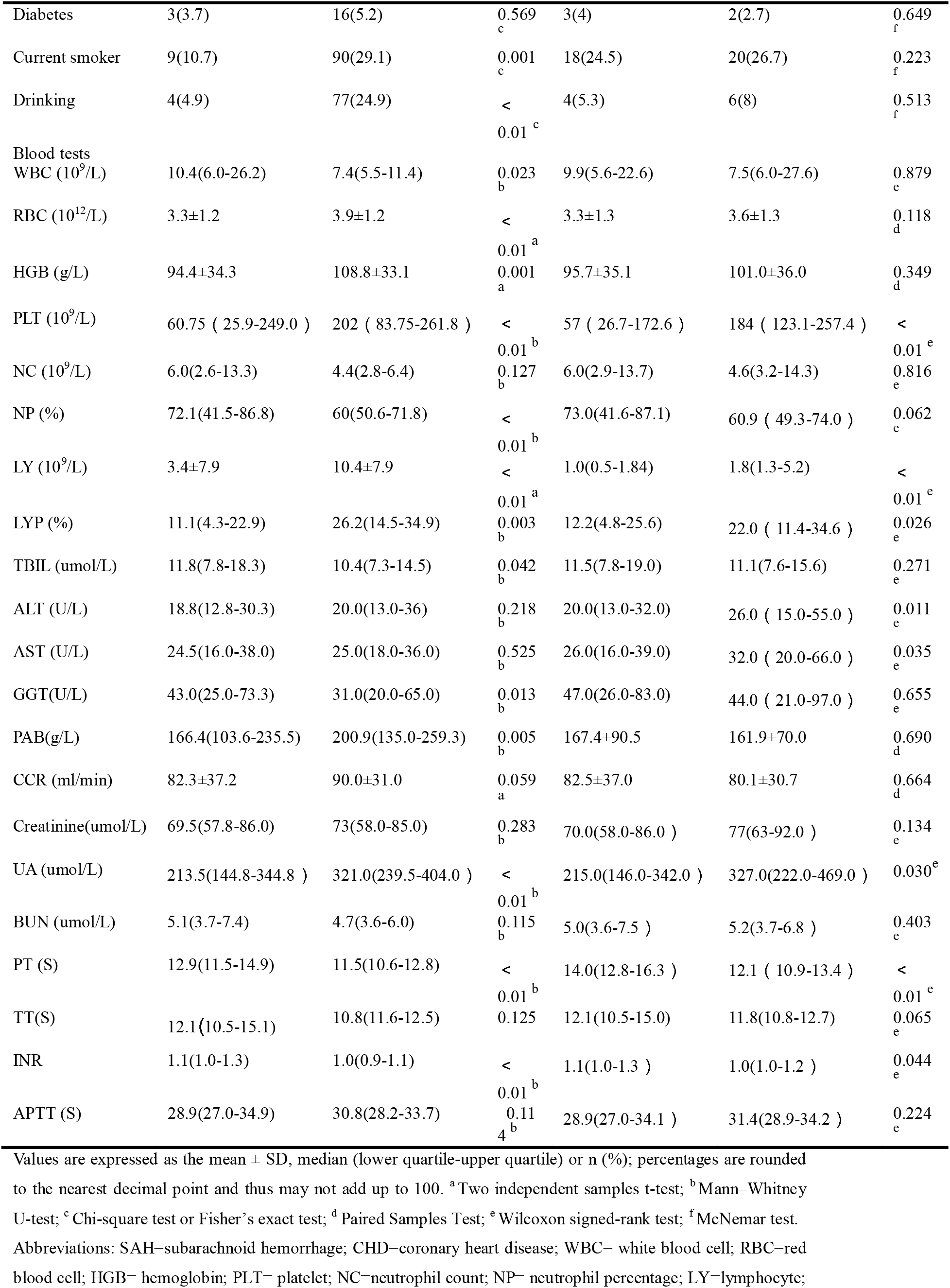
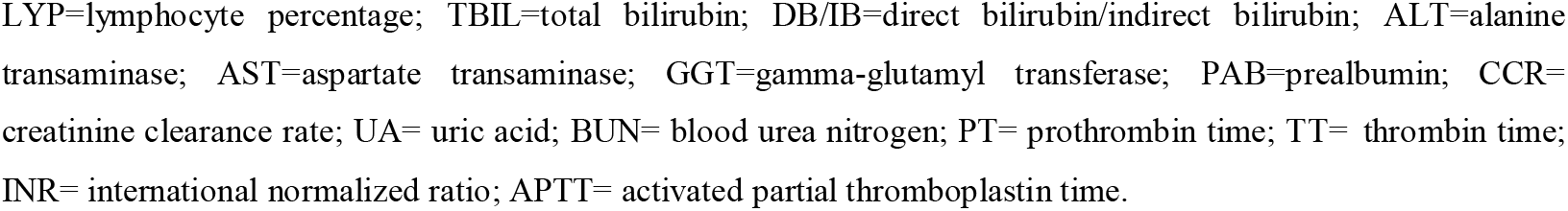
Patients’ profiles

| Variable | Before propensity score matching |  | p Value | After propensity score matching |  | p Value |
| --- | --- | --- | --- | --- | --- | --- |
|  | Active cancer with SAH (n=82) | Active Cancer alone (n=309) |  | Active cancer with SAH (n=75) | Active Cancer alone (n=75) |  |
| Age (year) | 53.0(44.0-66.0) | 52(40.0-63.0) | 0.450<sub>b</sub> | 52.3±17.2 | 56.3±15.6 | 0.146<sub>d</sub> |
| Male (%) | 56(68.3) | 193(62.4) | 0.049<sub>c</sub> | 51(69.3) | 46(61.3) | 0.303<sub>f</sub> |
| Previous strokes | 8(9.7) | 5(1.6) | < 0.01<sup>c</sup> | 1(1.3) | 2(2.7) | 0.560<sub>f</sub> |
| CHD | 2(2.4) | 4(1.3) | 0.454<sub>c</sub> | 2(2.7) | 1(1.3) | 0.560<sub>f</sub> |
| Hypertension | 17(20.7) | 39(12.6) | 0.050<sup>c</sup> | 16(21.3) | 10(13.3) | 0.196<sub>f</sub> |
| Diabetes | 3(3.7) | 16(5.2) | 0.569<sub>c</sub> | 3(4) | 2(2.7) | 0.649<sub>f</sub> |
| Current smoker | 9(10.7) | 90(29.1) | 0.001<sub>c</sub> | 18(24.5) | 20(26.7) | 0.223<sub>f</sub> |
| Drinking | 4(4.9) | 77(24.9) | < 0.01<sub>c</sub> | 4(5.3) | 6(8) | 0.513<sub>f</sub> |
| Blood tests |  |  |  |  |  |  |
| WBC (10<sup>9</sup>/L) | 10.4(6.0-26.2) | 7.4(5.5-11.4) | 0.023<sub>b</sub> | 9.9(5.6-22.6) | 7.5(6.0-27.6) | 0.879<sub>e</sub> |
| RBC (10<sup>12</sup>/L) | 3.3±1.2 | 3.9±1.2 | < 0.01<sub>a</sub> | 3.3±1.3 | 3.6±1.3 | 0.118<sub>d</sub> |
| HGB (g/L) | 94.4±34.3 | 108.8±33.1 | 0.001<sub>a</sub> | 95.7±35.1 | 101.0±36.0 | 0.349<sub>d</sub> |
| PLT (10<sup>9</sup>/L) | 60.75 ( 25.9-249.0 ) | 202 ( 83.75-261.8 ) | < 0.01<sub>b</sub> | 57 ( 26.7-172.6 ) | 184 ( 123.1-257.4 ) | < 0.01<sub>e</sub> |
| NC (10<sup>9</sup>/L) | 6.0(2.6-13.3) | 4.4(2.8-6.4) | 0.127<sub>b</sub> | 6.0(2.9-13.7) | 4.6(3.2-14.3) | 0.816<sub>e</sub> |
| NP (%) | 72.1(41.5-86.8) | 60(50.6-71.8) | < 0.01<sub>b</sub> | 73.0(41.6-87.1) | 60.9 ( 49.3-74.0 ) | 0.062<sub>e</sub> |
| LY (10<sup>9</sup>/L) | 3.4±7.9 | 10.4±7.9 | < 0.01<sub>a</sub> | 1.0(0.5-1.84) | 1.8(1.3-5.2) | < 0.01<sub>e</sub> |
| LYP (%) | 11.1(4.3-22.9) | 26.2(14.5-34.9) | 0.003<sub>b</sub> | 12.2(4.8-25.6) | 22.0 ( 11.4-34.6 ) | 0.026<sub>e</sub> |
| TBIL (umol/L) | 11.8(7.8-18.3) | 10.4(7.3-14.5) | 0.042<sub>b</sub> | 11.5(7.8-19.0) | 11.1(7.6-15.6) | 0.271<sub>e</sub> |
| ALT (U/L) | 18.8(12.8-30.3) | 20.0(13.0-36) | 0.218<sub>b</sub> | 20.0(13.0-32.0) | 26.0 ( 15.0-55.0 ) | 0.011<sub>e</sub> |
| AST (U/L) | 24.5(16.0-38.0) | 25.0(18.0-36.0) | 0.525<sub>b</sub> | 26.0(16.0-39.0) | 32.0 ( 20.0-66.0 ) | 0.035<sub>e</sub> |
| GGT(U/L) | 43.0(25.0-73.3) | 31.0(20.0-65.0) | 0.013<sub>b</sub> | 47.0(26.0-83.0) | 44.0 ( 21.0-97.0 ) | 0.655<sub>e</sub> |
| PAB(g/L) | 166.4(103.6-235.5) | 200.9(135.0-259.3) | 0.005<sub>b</sub> | 167.4±90.5 | 161.9±70.0 | 0.690<sub>d</sub> |
| CCR (ml/min) | 82.3±37.2 | 90.0±31.0 | 0.059<sub>a</sub> | 82.5±37.0 | 80.1±30.7 | 0.664<sub>d</sub> |
| Creatinine(umol/L) | 69.5(57.8-86.0) | 73(58.0-85.0) | 0.283<sub>b</sub> | 70.0(58.0-86.0 ) | 77(63-92.0 ) | 0.134<sub>e</sub> |
| UA (umol/L) | 213.5(144.8-344.8 ) | 321.0(239.5-404.0 ) | < 0.01<sub>b</sub> | 215.0(146.0-342.0 ) | 327.0(222.0-469.0 ) | 0.030<sub>e</sub> |
| BUN (umol/L) | 5.1(3.7-7.4) | 4.7(3.6-6.0) | 0.115<sub>b</sub> | 5.0(3.6-7.5 ) | 5.2(3.7-6.8 ) | 0.403<sub>e</sub> |
| PT (S) | 12.9(11.5-14.9) | 11.5(10.6-12.8) | < 0.01<sub>b</sub> | 14.0(12.8-16.3 ) | 12.1 ( 10.9-13.4 ) | < 0.01<sub>e</sub> |
| TT(S) | 12.1(10.5-15.1) | 10.8(11.6-12.5) | 0.125<sub>b</sub> | 12.1(10.5-15.0) | 11.8(10.8-12.7) | 0.065<sub>e</sub> |
| INR | 1.1(1.0-1.3) | 1.0(0.9-1.1) | < 0.01<sub>b</sub> | 1.1(1.0-1.3 ) | 1.0(1.0-1.2 ) | 0.044<sub>e</sub> |
| APTT (S) | 28.9(27.0-34.9) | 30.8(28.2-33.7) | 0.114<sub>b</sub> | 28.9(27.0-34.1 ) | 31.4(28.9-34.2 ) | 0.224<sub>e</sub> |

Values are expressed as the mean ± SD, median (lower quartile-upper quartile) or n (%); percentages are rounded to the nearest decimal point and thus may not add up to 100. <sup>a</sup>Two independent samples t-test; <sup>b</sup>Mann–Whitney U-test; <sup>c</sup>Chi-square test or Fisher’s exact test; <sup>d</sup>Paired Samples Test; <sup>e</sup>Wilcoxon signed-rank test; <sup>f</sup>McNemar test. Abbreviations: SAH=subarachnoid hemorrhage; CHD=coronary heart disease; WBC= white blood cell; RBC=red blood cell; HGB= hemoglobin; PLT= platelet; NC=neutrophil count; NP= neutrophil percentage; LY=lymphocyte;
LYP=lymphocyte percentage; TBIL=total bilirubin; DB/IB=direct bilirubin/indirect bilirubin; ALT=alanine transaminase; AST=aspartate transaminase; GGT=gamma-glutamyl transferase; PAB=prealbumin; CCR= creatinine clearance rate; UA= uric acid; BUN= blood urea nitrogen; PT= prothrombin time; TT= thrombin time; INR= international normalized ratio; APTT= activated partial thromboplastin time.

A multivariate logistic regression analysis demonstrated that two variables were significantly relating to the onset of SAH: having previous stroke (*p*=0.006; odds ratio [OR], 3.85; 95% CI, 1.73 to 5.06) and prothrombin time (*p*=0.028; odds ratio [OR], 1.14; 95% CI, 1.02 to 1.29) (Appendix 11: Table S7).

Nevertheless, the results of a univariate analysis of the propensity score-matched groups revealed that eight variables were significantly related to the onset of SAH: the level of platelet, lymphocyte, lymphocyte percentage, alanine transaminase, aspartate transaminase, uric acid, prothrombin time, and international normalized ratio (Table 3). Multivariate analysis revealed that only two variable was significantly related to the onset of SAH: the level of platelet (*p*<0.001; odds ratio [OR], 0.87; 95% CI, 0.69 to 0.98); prothrombin time (*p*<0.001; odds ratio [OR], 1.559; 95% CI, 1.229 to 1.976) (Appendix 4:Figure S4, Appendix 8:Table S12).

### Receiver operating characteristic curves

The receiver operating characteristic (ROC) curves for identifying the onset of SAH in patients with active cancer from prothrombin time, platelet, and joint predictor (combing prothrombin time and platelet) were shown in Figure 8. Using prothrombin time and platelets as covariates X_1_ and X_2_ respectively, the joint predictive factor expression is obtained: Logit(P)=β_1_/β_1_*X_1_+β_2_/β_1_*X_2_ (Appendix 6). Receiver operating characteristic curve of the joint predictive factor showed the largest AUC of 0.8131 (95% CI 0.745 to 0.881, *P*<0.001), indicating good overall accuracy of the test. The optimum diagnostic cut-off value for the joint predictive factor calculated from the ROC was 11.719, with a sensitivity of 65.3% and specificity of 89.3% (Figure 4, Appendix 9: Table S13).

**Figure 4.** Receiver operating characteristic curves obtained with different discriminatory models for predicting the onset of SAH in patients with active cancer. The joint predictive factor showed the largest AUC of 0.8131 (95% CI 0.745 to 0.881, P<0.001). The optimum cut-off point was 11.719. At this cut-off value, the sensitivity was 65.3% and the specificity was 89.3%. CI: Confidence interval; ROC: Receiver operator characteristic. AUC: Area under the curve.

## DISCUSSION

Subarachnoid hemorrhage in patients with active cancer has received insufficient attention^8, 9^. The clinical features of SAH in patients with active cancer are noticeable^10^, and may be an initial feature of the malignancy, complication of active malignancy or delayed consequence of cancer and its management ^11^. In the present study, most SAH in patients with active cancer had common clinical manifestations, such as headache, nausea and vomiting which correlated with the findings of the previous studies ^6, 7, 12, 13^. Further, this study represented the largest clinical series of SAH in patients with cancer. It was found that most SAH in patients with active cancer developed SAH within the first 6 months after diagnosis of cancer, suggesting that as soon as cancer diagnosis is established, measures should be taken to prevent SAH. Besides, some cancer patients have SAH as the first presentation. Subarachnoid hemorrhage had been reported in 1956 as the first presentation ^14^, which indicate that when the pathogenesis of SAH was unexplained, the measures should be taken to determine the causes including aneurysms negative in first examination and occult cancer ^15, 16^.

On the other hand, the intracranial hemorrhage in patients with cancer were considered catastrophic and terminal event^2^. As a result, the clinical feature of cerebral hemorrhage and subarachnoid hemorrhage in patients with cancer remained not fully understood. In the present research, the severity of SAH was assessed with Hunt & Hess grade. It was revealed that nearly one in six patients have severe Hunt & Hess grade score. Among them, liquid tumors were rated as the most severe Hunt & Hess grade, which corroborates with the findings of two previous studies^17, 18^. This suggests that SAH in patients with cancer is serious and requires prompt and aggressive rescue measures. For the sake of exploring 30 days and 90 days mortality of SAH patients with active cancer, Kaplan–Meier curves based on cancer type and etiology were established. Subsequently, it was found that SAH patients with active cancer had poor outcomes, causing thirty-day mortality of 41.5%, and ninety-day mortality 52.0%. Moreover, it was also found that tumor types and the causes of SAH were related to the median survival time of SAH in patients with cancer, which was also similar to previous studies^6^. Nevertheless, the present study added new insight about SAH in patients with active cancer.

Intracranial hemorrhage, including SAH in patients with cancer have been found with poor prognosis^6, 7^. However, predictors of mortality of SAH in patients with active cancer have not been investigated. Here, cox regression models were configured to explore the predictors of mortality. It was that liquid tumor, activated partial thromboplastin time, prealbumin, coagulopathy, gamma-glutamyl transferase, atrial fibrillation, and cryptogenic subarachnoid hemorrhage were independent factors of survival in SAH patients with active cancer. This provided useful information facilitating clinicians to take up appropriate therapeutic measures.

Further, univariate analysis and multivariate analysis were also applied to reveal the independent risk factors of occurrence of SAH in patients with active cancer. It was noted that previous stroke, prothrombin time were the independent risk factors of occurrence of SAH in patients with active cancer. Additionally, to reveal the independent risk for cancer related SAH, a propensity score matching (PSM) eliminating the impacts of conventional vascular risks on SAH in cancer patients was performed. The PSM found that the decrease of platelet and prolonged PT time were the independent risk factors of cancer related SAH. In 2014, two studies provided a link between thrombocytopenia and prognosis or development of stroke (ischemic or hemorrhagic) ^19, 20^. Therefore, the present study confirmed that decrease of platelet and prolonged PT associated with cancer related SAH.

However, decreased platelet and prolonged PT are also found in cancers and other diseases as a common coagulation marker ^19, 21, 22^. Considering that development of SAH in active cancer patients may be caused by the combined effects of the decrease of platelet and prolonged PT, the joint predictive factor was calculated. That the area under the ROC curve of the joint predictive factor was larger than that of each decreased platelet and prolonged PT and the sensitivity and specificity of the joint predictive factor were highest. This suggests that that the joint predictive factor could serve as a potential biomarker of cancer related SAH. Furthermore, by having the cutoff value of the joint predictive factor equal to 11.719, clinicians can discover the patient at high risk of SAH for cancer patients and identify the cancer related SAH from other etiologic SAH.

This study had some limitations which should also be mentioned before generalizing the present findings. Generally, this study was a retrospective comparison with propensity score-matching to minimize the bias in patient selection, but unobserved confounders remained. On the other hand, further research is needed because previous studies have suggested that factors affecting the occurrence of SAH include histological characteristics of the tumor, tumorous position, cancerous aneurysm and direct invasion of meningeal blood vessels, selective serotonin reuptake inhibitor as well as estrogen replacement therapy^6, 23-30^.

In conclusion, cancer patients with SAH often have poor prognosis. The decrease in platelet and prolonged PT are the independent risk factor of cancer related SAH. Further, the joint predictive factor with cutoff value equal to 11.719 should serves as a novel biomarker to facilitate clinicians in establish the patients at high risk of SAH and identify cancer related SAH from other etiologic SAH. However, further studies should be conducted to confirm the present findings.

